# Combined in vitro differentiation and cell sorting-based isolation of highly pure mouse bone marrow-derived basophils

**DOI:** 10.1101/2024.04.25.591228

**Authors:** Adriana Rosa Gambardella, Valentina Tirelli, Sara Andreone, Jacopo Mancini, Fabrizio Mattei, Giovanna Schiavoni

**Affiliations:** Department of Oncology and Molecular Medicine, Istituto Superiore di Sanità, Viale Regina Elena, 299, 00161, Rome, Italy; Core Facilities, Istituto Superiore di Sanità, Viale Regina Elena, 299, 00161, Rome, Italy

## Abstract

Basophils constitute a rare population of granulocytes with key functions in allergies, immunodeficiencies and cancer. The scarcity of basophils in the human blood and tissues constitutes a considerable limit for the study of these cells. Interleukin-3 (IL-3) is a cytokine that stimulates both the differentiation and the expansion of basophils from bone marrow (BM) precursors by positively regulating the expression of the IL3Ra receptor. We have standardized an *in vitro* differentiation protocol of mouse basophils (mBaso) from BM precursors through culture in presence of IL-3 for 10 days followed by cell sorting. At the end of the 10-day differentiation, a considerable number of mBaso can be obtained and cell sorting procedures further improved the isolation of an extraordinarily pure (>98%) and vital FcεR1^+^ CD11c^-^ c-kit^-^ mBaso population. Phenotypic analysis revealed that terminally differentiated (day 10) unsorted mBaso cultured for 24 hours showed a decrease in basophilic lineage (c-kit^-^) and an increase of mastocytic lineage (c-kit^+^) and reduced the expression of basophil markers FcεRI, CD49b and CD200R either in absence of stimuli or following activation with the alarmin IL-33, indicating cell dedifferentiation. In contrast, terminally differentiated and FcεR1^+^ CD11c^-^ c-kit^-^ sorted mBaso do not dedifferentiate in mast cells when placed in culture, and responded to IL-33 stimulation by up-regulating the activation marker CD63 without down-modulation of FcεRI and CD200R3. These evidences highlight that *in vitro* differentiation followed by cell sorting is a useful method to obtain elevated numbers of highly pure mBaso that preserve their lineage markers and thus are suitable for conducting the desired functional studies.

## 1) Introduction

Although basophils account only for <1% of blood leukocytes (Gauvreau et al., 2005; Borriello et al., 2014), they carry out central roles in clearing pathogens (Galli, 2000; Siracusa et al., 2013), and initiating inflammatory responses in allergy, autoimmunity, fibrosis, COVID-19 and cancer (Miyake et al., 2020; Sharma et al., 2015; Murdaca et al., 2021; Schiechl et al., 2016; Hayes et al., 2020; Marone et al., 2020). As other granulocytes, basophils originate from hematopoietic immature cells and mature into terminally differentiated cells in the bone marrow (BM). Basophils are released in blood vessels and circulate in the peripheral blood (Yamanishi et al., 2017). Circulating basophils have a short life-span that account for 2,5 days in mice and their replacement is constantly resupplied from BM to the blood (Arinobu et al., 2005; Poto et al., 2022). During inflammation, basophils can extravasate from the bloodstream and infiltrate damaged tissues where they were activated by the secretion of inflamed tissue-released immunomodulatory cytokines, such as interleukin-25 (IL-25), IL-33 and thymic stromal lymphopoietin (TSLP) (Siracusa et al., 2013; Ohnmacht et al., 2008). Basophil activation results in the release of the type 2 cytokines IL-4, IL-5, IL-9 and IL-13 (Gambardella et al., 2022; Peng et Siracusa, 2021). Since the number of circulating basophils may be altered under certain pathologic conditions they can represent prognostic markers of disease, such as of survival outcomes in tumors (Liu et al., 2020).

Despite the relevance of their immune functions, the scarcity of circulating human basophils (hBaso) and the difficulties in obtaining these cells from human tissues *ex vivo* represent the main challenges in conducting studies to investigate the role of these rare immune cells in health and disease. BM-derived mouse basophils (mBaso) differentiated *in vitro* represent a model system with good similarities with the human cells. As already reported, mBaso and hBaso may differentiate *in vitro* by culturing mouse BM-derived cells or human CD34^+^ cells in presence of IL-3 for 10 days (Gambardella et al., 2022; Valent et al., 1989). IL-3 stimulates both the expansion and the differentiation of either mBaso and hBaso by upregulating *Il3ra* mRNA and in turn IL-3 receptor expression (Li et al., 2022).

Here, we illustrate a methodology to obtain a considerable number of highly pure mBaso by culturing mouse BM-derived cells in presence of IL-3 for 10 days followed by cell sorting-based strategies. Interestingly, both mBaso and mast cells share morphological similarity, such as the expression of full tetrameric form of the high affinity receptor FcϵRI and the storage of histamine in cytoplasmic granules and its release, together with other proinflammatory mediators (Marone et al., 2020; Yamanishi et al., 2017). After 10 days of differentiation in the presence of IL-3, mBaso are FACS-sorted as FcεR1^+^ CD11c^-^ c-kit^-^ to remove contaminating mast cells (FcεR1^+^ c-kit^+^) and dendritic cells (CD11c^+^), yielding more than 98% pure population (Gambardella et al., 2022). As an added value, the purity of the obtained mBaso population is reported by flow cytometry phenotypic analysis. Through these standardized in vitro differentiation and cell sorting-based methodology, we generated a significant number (~ 8-10×10^6^) of highly pure mBaso (FcεR1^+^ CD11c^-^ c-kit^-^) suitable to perform desired functional studies.

## 2) Materials

### Animals

- 6-10 week old C57Bl/6 mice (Charles River Laboratories, Calco, Italy)

### Disposables

- 75 cm^2^ screw cap with filter cell culture flasks (#ET7076, Euroclone)
- 50 mL conical-bottom centrifuge tubes (#ET5050B, Euroclone)
- 5 mL polystirene round bottom tubes (#352054, Corning)
- 1 mL syringe (#303175, BD Plastipack)
- 100 μm filters (Filcon sterile syringe-type; #340606, BD Biosciences)

### Cells and reagents

- Bone morrow cells isolated from the aforementioned mice
- Phosphate-buffered saline (PBS; #ECB4004L, EuroClone)
- MACS buffer (1% FBS, 2 mM EDTA in PBS)
- FACS buffer (2% FBS, 0,1% Sodium Azide in PBS)
- Recombinant mouse (rm)IL-3 (#403-ML, R&D System)
- rmIL-33 (#580504, Biolegend)
- RPMI 1640 (#ECB9006L, EuroClone)
- Fetal Bovine Serum (FBS; #ECS5000L, EuroClone)
- Antibiotic-Antimycotic (#ECM0010D, EuroClone)
- Glutamine (#ECB3000D, EuroClone)
- Sodium Pyruvate (#ECM0542D, EuroClone)
- RPMI 1640 culture medium containing 10% FBS, 1% Antibiotic-Antimycotic, 1% Glutamine (namely RPMI complete medium)
- Ammonium-chloride potassium (ACK) red blood cell lysis buffer (See *Note 2*)
- RPMI 1640 supplemented with 1% of sodium pyruvate (namely mBaso medium)
- Sytox Blue Dead Cell Stain (#S34857, Thermo Fisher Scientific)

### Antibodies

- FITC anti-mouse FceR1 (clone MAR-1, Biolegend)
- PE-Cy7 anti-mouse CD117/c-kit (clone 2B8, Biolegend)
- APC anti-mouse CD11c (clone N418, Biolegend)
- PE anti-mouse CD49b (clone DX5, Miltenyi Biotec) or
- PE anti-mouse CD63 (clone NVG-2, Biolegend)
- APC anti-mouse CD200R3 (clone Ba13, Biolegend)

### Equipment

- Class II Laboratory Biosafety Cabinet VBH 75 MP (#8071, Steril)
- Humidified cell culture incubator at +37 °C with 5% CO_2_ (Thermo Fisher Scientific)
- Bench top centrifuge (SL Plus Series, Thermo Fisher Scientific)
- Gallios Flow Cytometer (Beckman Coulter)
- MoFlo Astrios EQ Cell Sorter (Beckman Coulter)

### Software

- Kaluza Analysis Software (Beckman Coulter)
- Summit Software System for cell sorting analysis (Cytomation)

## 3) Methods

### 3.1 Extraction of bone marrow from mouse tibias and femurs

1. Sacrifice C57Bl/6 mice to collect tibias and femurs.
2. Strip all tissue residues from bones by the use of small and sterile pliers and scissors (see *Note 1*).
3. Extract BM from both tibias and femurs by using 1 mL syringe needle. To do this, inject RPMI complete medium (see *Section 2, Cells and reagents*) in the bone central canal to allow the leakage of BM from bones.
4. Harvest BM in 50 mL centrifuge tube and resuspend in cold RPMI complete medium.
5. Disaggregate BM by carefully pipetting to obtain a cell suspension.
6. Once completed BM cell disaggregation, centrifuge BM cell suspension at 1500 rpm, 4°C, 5 minutes to eliminate cell debris. Following centrifugation, discard the supernatant.
7. Resuspend the pellet with a manual slight swinging of the centrifuge tube.
8. Wash the pellet by adding 10-15 mL pre-warmed PBS and by centrifuging 1500 rpm, 4°C, 5 minutes. To discard PBS, aspirate supernatant by employing a 10 mL serological pipette.
9. Subsequently, resuspend the pellet in 4 mL pre-warmed ACK red blood cell lysis buffer (ACK; see *Note 2*) for 4 minutes at room temperature (RT) to eliminate erythrocytes. Mix, gently but frequently, BM cells incubated with ACK to ensure an uniform cell resuspension and, therefore, an efficient red blood cell lysis.
10. At the end of incubation, dilute BM cells immediately by adding 15-20 mL cold RPMI complete medium.
11. Wash cell suspension by centrifuging 1500 rpm, 4°C, 5 minutes. Discard the supernatant.

### 3.2 In vitro differentiation of mouse bone marrow -derived basophils

1. Plate 30 × 10^6^ BM-derived cells in a 75 cm^2^ cell culture flask in 20 mL of mBaso medium (see *Cells and reagents Section*) plus 2 ng/mL of rmIL-3. Add rmIL-3 directly in the culture medium.
2. At day 4 and 7 of differentiation, harvest non-adherent cells by aspirating the culture supernatant with a serological pipette.
3. Transfer the culture supernatant, that contains differentiating mBaso, in a 50 mL centrifuge tube.
4. Pellet cells by centrifuging 1500 rpm, 4°C, 5 minutes.
5. Following centrifugation, discard supernatant and resuspend cells accurately in 20 mL of fresh mBaso medium. Therefore, re-seed cell suspension into a new 75 cm^2^ cell culture flask in presence of 2 ng/ml of rmIL-3. (The time schedule of in vitro differentiation of mBaso is illustrated in *Figure 1*).
6. At day 10, mBaso are terminally differentiated. Collect non-adherent mBaso by aspirating cell culture supernatant.
7. Transfer cell culture supernatant in a 50 mL centrifuge tube.
8. Pellet mBaso by centrifuging 1500 rpm, 4°C, 5 minutes. Following centrifugation, discard the supernatant.
9. Resuspend mBaso in the mBaso medium (for cytokine stimulation) or MACS/FACS buffer (for sorting procedures/cell staining and flow cytometry analysis) depending on specific experimental requirements.

**Figure 1.**
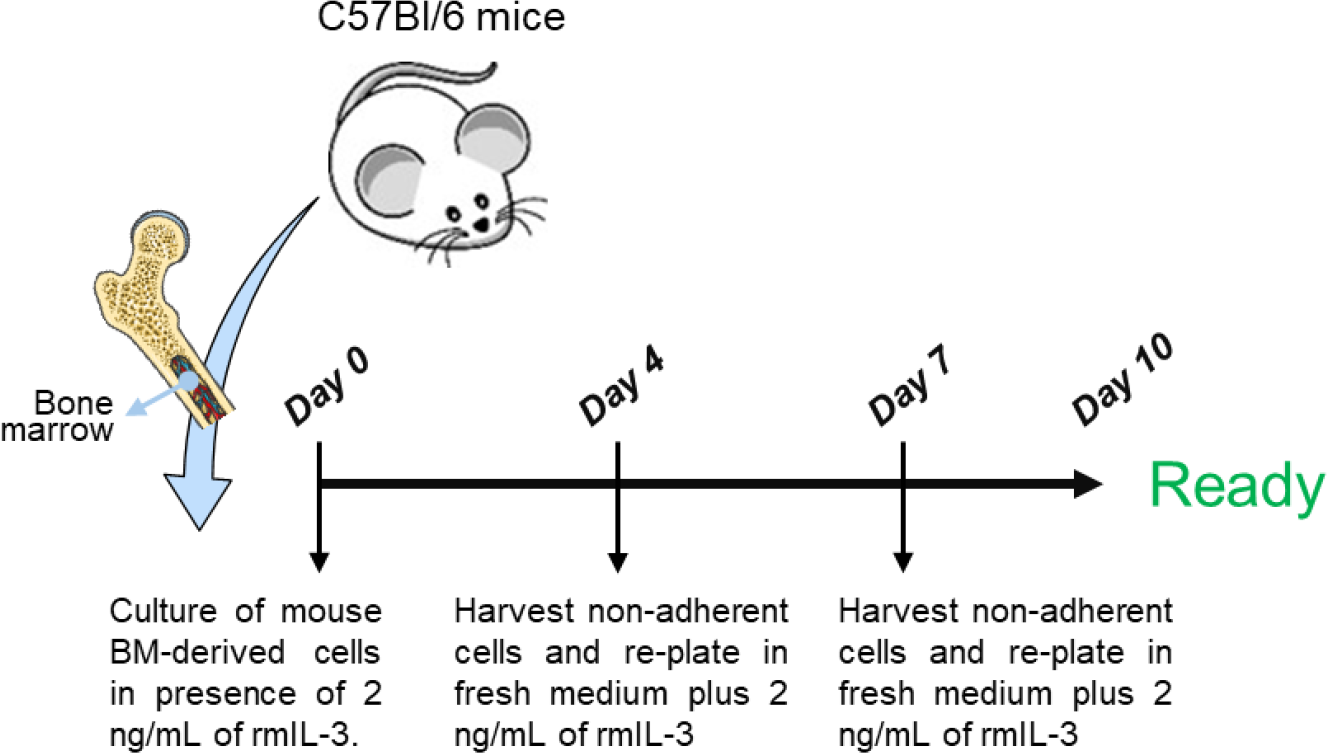
Time schedule of in vitro differentiation protocol of mouse basophils (mBaso). Once bone marrow (BM)-derived cells are obtained from mice, cells are cultured with 2 ng/mL of recombinant mouse (rm)IL-3 and refreshed each 3-4 days. At day 10, mBaso are terminally differentiated.

### 3.3 Staining of unsorted day 10-differentiated mouse basophils

1. At day 10, once collected the terminally differentiated non-adherent mBaso by aspirating the culture supernatant and pelleted as above described, wash cells twice by bringing up to volume until 40-45 mL with cold FACS buffer and by centrifuging 1500 rpm, 4°C, 5 minutes. Discard the supernatant.
2. Stain mBaso within a 5 mL polystirene round bottom tube with the following monoclonal antibodies: FITC anti-mouse FcεR1, PE-Cy7 anti-mouse CD117/c-kit, PE anti-mouse CD49b, APC anti-mouse CD200R3 (see *Note 3* for staining conditions; *Figure 2, “day 10 mBaso”)*.

**Figure 2.**
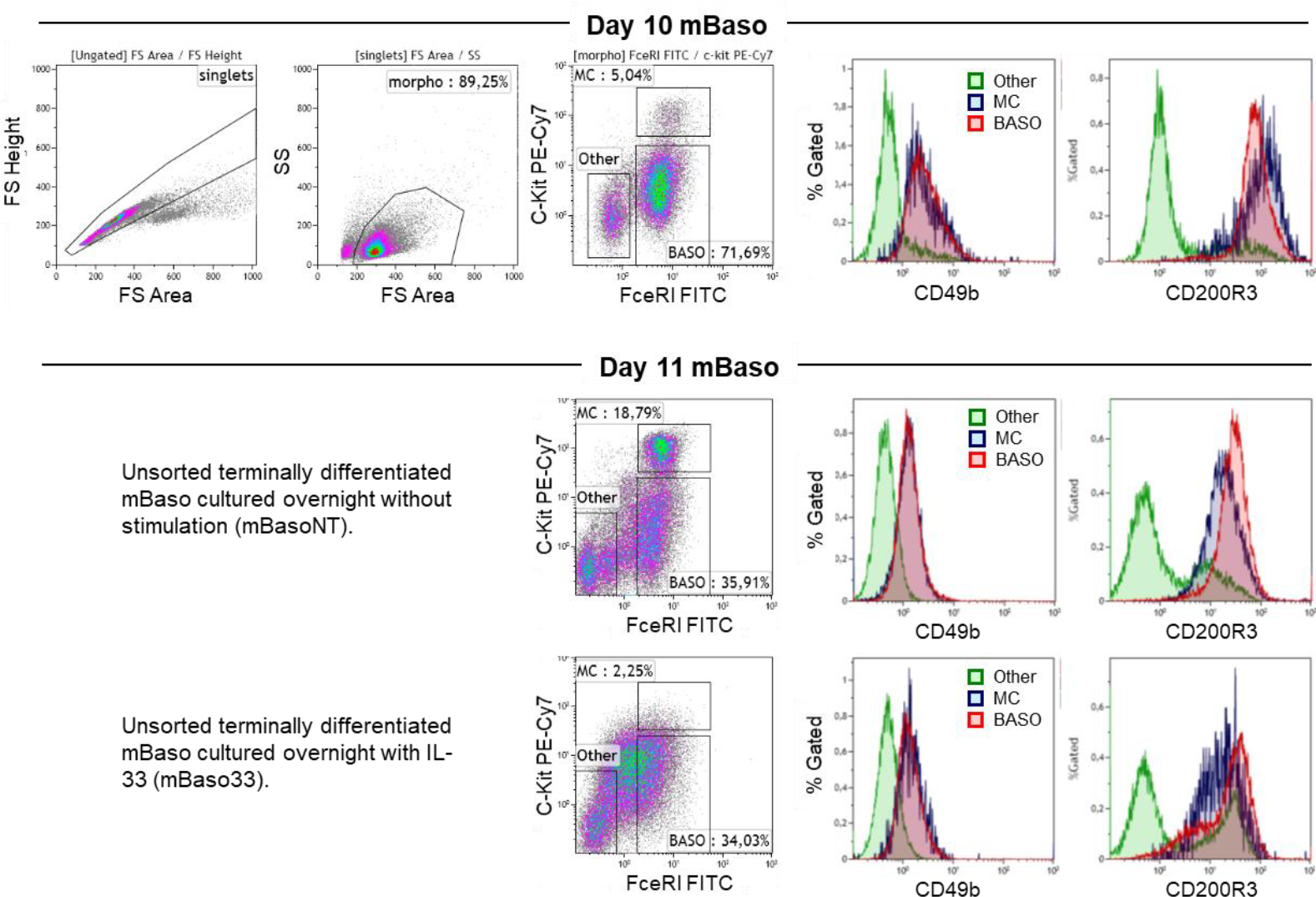
Phenotyipic analysis of mouse basophils differentiated in vitro with IL-3. Bone marrow (BM)-derived cells are cultured with 2 ng/mL recombinant mouse (rm)IL-3 for 10 days. At day 10, cells are harvested and stained with the following monoclonal antibodies FITC anti-mouse FcεR1, PE-Cy7 anti-mouse CD117/c-kit, PE anti-mouse CD49b, APC anti-mouse CD200R3. Flow cytometry analysis reveal the yields around 70% FcεR1+ c-kit-mBaso and only 5% FcεR1+ c-kit+ mast cells (MC). Both mBaso and MC express FcεR1, CD49b and CD200R3, while they can be distinguished by c-kit expression. At day 10, terminally differentiated mBaso are left unstimulated (mBasoNT) or stimulated overnight with 100 ng/mL rmIL-33 (mBaso33). At day 11, flow cytometry analysis reveal that upon overnight stimulation with 100 ng/mL of rmIL33, mBaso33 down-regulate FcεRI, CD49b and CD200R3. Moreover, following overnight culture of mBasoNT, mBaso percentage decreases while increasing percentage of MC is observed.

### 3.4 Cytokine stimulation of unsorted mouse basophils

1. Resuspend terminally differentiated and unsorted mBaso (day 10) at a concentration of 1×10^6^ cells/mL in mBaso medium.
2. Seed 1×10^6^ mBaso into 48 multiwell plate in a final volume 1 mL.
3. Left unstimulated the control cell cultures (mBasoNT) and stimulate with 100 ng/mL rmIL-33 the other mBaso cultures (mBaso33). Incubate mBasoNT and mBaso33 cultures overnight (until day 11) at 37°C (see *Note 4*).

### 3.5 Staining of unsorted mouse basophils following overnight culture (day 11)

1. Following overnight culture (day 11; see *Section 3*.*4*), harvest mBasoNT and mBaso33 by gently pipetting prior to transfer cells in a 5 mL polystirene round bottom tube (*See Note 5*).
2. Wash cells twice by bringing up to volume until 4-4,5 mL with FACS buffer and by centrifuging 1500 rpm, 4°C, 5 minutes. Discard the supernatant.
3. Stain mBaso in the 5 mL polystirene round bottom tube, in which they are already contained, with the following monoclonal antibodies: FITC anti-mouse FcεR1, PE-Cy7 anti-mouse CD117/c-kit, PE anti-mouse CD49b, APC anti-mouse CD200R3 (see *Note 3* for staining conditions).

### 3.6 Phenotypic analysis of unsorted day 10 and day 11 mBaso

1. Flow cytometry of day 10-differentiated and day 11 (overnight cultured) unsorted mBaso is performed on Gallios Flow Cytometer equipped with 488 nm (blue), 638 nm (red), and 405 nm (violet) lasers.
2. To acquire data, the following channels are used: SSC, FSC, FL1 (488 nm laser, 525/20 BP), FL2 (488 nm laser, 575/20 BP), FL5 (488 nm laser, 755LP BP), and FL6 (638 nm laser, 660/20 BP).
3. Laser voltage (gain) and compensation are set using unstained cells and cells stained with each fluorochrome separately.
4. At least 10,000 events with normal SSC and FSC are acquired per sample, ideally at a flow rate < 500 events/second.
5. For analysis, in the Kaluza software generate dot plot masks based on the FSC/SSC parameters and all fluorescence channel combinations to adjust compensation as necessary.
6. Exclude debris by selecting single cells (singlets) using FSCA/FSCH parameters. Then, select cells with normal morphology (morpho) using FSC/SSC mask gated on “singlets” (*Figure 2, “Day 10 mBaso”*).
7. From the mask FcεR1 FITC/c-kit PE gated in “morpho” generate regions to identify basophils (BASO) as FcεR1^+^c-kit^-^, mast cells (MC) as FcεR1^+^c-kit^+^, and other cells (mainly dendritic cells, see below) as FcεR1^-^c-kit^-^ and obtain the relative percentage of each gated population by selecting “% of Gated”.
8. Obtain histogram plots for CD49b PE and CD200R3 APC and generate overlays by alternatively gating on the three murine subpopulations (Baso, MC and others; see *Figure 2, “Day 10 mBaso” and “Day 11 mBaso”*) to check expression intensity of the two markers in the three cell populations.

### 3.7 Sorting of terminally differentiated mouse basophils

1. At day 10, collect the terminally differentiated non-adherent mBaso by aspirating the culture supernatant as above described (*Section 3*.*2*).
2. To carry out cell sorting procedures, wash mBaso with 10 mL MACS buffer (*Section 2, Cells and reagents*) by centrifuging 1500 rpm, 4°C, 5 minutes to ensure the elimination of medium and serum residues which may adversely affect flow cytometry workflow.
3. Resuspend pellet in 3 mL of cold MACS buffer prior to transfer all cells (~ 25-30×10^6^) in a 5 mL polystirene round bottom tube for staining.
4. Centrifuge to pellet mBaso 1500 rpm, 4°C, 5 minutes and discard supernatant.
5. Stain cells with the following monoclonal antibodies: FITC anti-mouse FcεR1, PE-Cy7 anti-mouse CD117/c-kit and APC anti-mouse CD11c (see *Note 6* for staining condition).
6. Wash labeled mBaso twice (1300 rpm, 4°C, 5 minutes) with 4 mL of cold MACS buffer.
7. Resuspend at a cellular concentration of 20-25×10^6^ mBaso/mL in cold MACS buffer (see *Note 7)*.
8. Before sorting, filter cell suspension through 100 μm filters (Filcon sterile syringe-type; see *Note 8*)
9. A highly pure population of mBaso (FcεR1+ CD11c-c-kit-) is sorted using the high-speed cell sorter MoFlo AstriosEQs, which is equipped with 4 laser lines (405 nm, 488 nm, 561nm and 642 nm) and 15 fluorescence parameters. Sorting is performed using a 100μm nozzle tip. The sheath pressure is maintained at approximately 30psi and the acquisition speed set at around 8000 events per second, to reduce the abort rate and increase sorting efficiency
10. The sorting strategy for mBaso (FcεR1^+^ CD11c^-^ c-kit^-^) is set as follows: first, dead cells are excluded based on Sytox blue positivity (see *Note 9*), and doublets are gated out using Forward scatter (FSC) Area versus side scatter (SSC) width. Then, CD11c^-^ and c-Kit^-^ cells are gated in, and finally, the FcεR1^+^ population is selected for sorting. The post-sorting analysis reveal a purity of mBaso > 98% (*Figure 3*).

**Figure 3.**
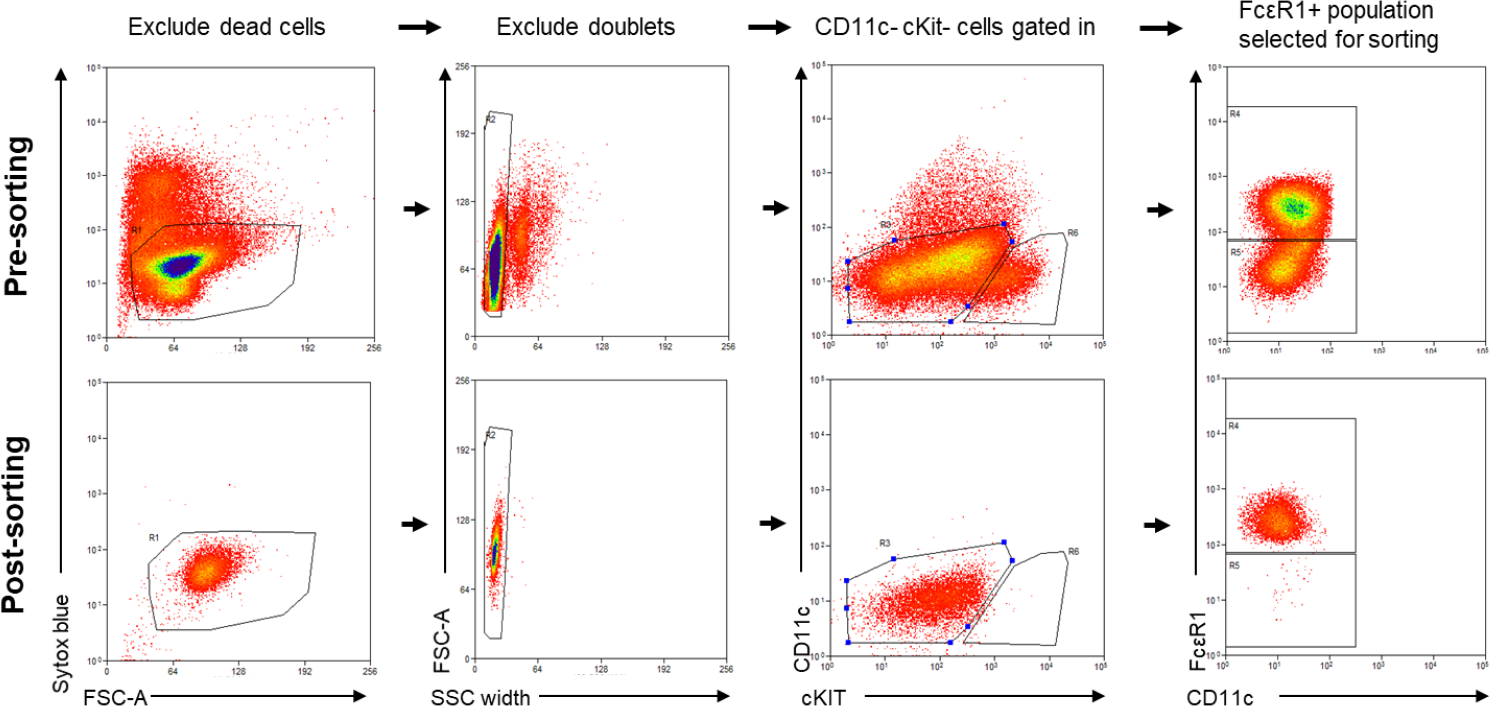
Cell sorting of FcεR1+ CD11c- c-kit- mouse basophils (mBaso). Terminally differentiated mBaso are stained with the following monoclonal antibodies: FITC anti-mouse FcεR1, PE-Cy7 anti-mouse CD117/c-kit and APC anti-mouse CD11c. Labeled mBaso are marked with Sytox blue and then run on cell sorter. Sytox blue positive dead cells are excluded. Doublets are left out using Forward scatter (FSC) Area versus side scatter (SSC) width. CD11c- c-kit- cells are gated in and, finally, FcεR1+ population is selected for sorting. Post-sorting analysis reveal a mBaso purity >98%.

### 3.8 Cytokine stimulation of sorted mouse basophils

1. Upon cell sorting, pellet FcεR1^+^ CD11c^-^ c-kit^-^ mBaso by centrifuging 1500 rpm, 4°C, 5 minutes. Discard supernatant.
2. Now, resuspend sorted mBaso at a concentration of 1×10^6^ cells/mL in mBaso medium.
3. Plate 1×10^6^ of sorted mBaso into 48 multiwell plate (final volume 1 mL).
4. Incubate cell culture at 37°C overnight (until day 11) unstimulated or stimulated with 100 ng/mL rmIL-33 (see *Note 4*).

### 3.9 Phenotypic analysis of sorted mBaso cultured overnight alone or with IL-33

1. Following overnight culture of sorted mBaso with IL-33 or left unstimulated (day 11; *See section 3*.*7*), harvest mBasoNT and mBaso33 by gently pipetting before they are transferred in a new 5 mL polystirene round bottom tube (*See Note 5*).
2. Wash mBaso twice (to ensure the elimination of medium and serum residues which may have a detrimental effect on flow cytometry workflow) by bringing up to volume until 4-4,5 mL with FACS buffer and by centrifuging 1500 rpm, 4°C, 5 minutes. Discard supernatant.
3. Stain cells with the following monoclonal antibodies: FITC anti-mouse FcεR1, PE-Cy7 anti-mouse CD117/c-kit, PE anti-mouse CD63, APC anti-mouse CD200R3 (see *Note 10* for staining conditions).
4. Wash labeled mBaso twice (1300 rpm, 4°C, 5 minutes) with FACS buffer.

### 3.10 Phenotypic analysis of sorted in vitro cultured mBaso

1. Flow cytometry of day 11 (overnight cultured) sorted mBaso is performed on Gallios Flow Cytometer (*See 3*.*6*).
2. To acquire data, the following channels are used: SSC, FSC, FL1 (488 nm laser, 525/20 BP), FL2 (488 nm laser, 575/20 BP), FL5 (488 nm laser, 755LP BP), and FL6 (638 nm laser, 660/20 BP).
3. Laser voltage (gain) and compensation are set using unstained cells and cells stained with each fluorochrome separately.
4. At least 10,000 events with normal SSC and FSC are acquired per sample, ideally at a flow rate < 500 events/second.
5. For analysis, in the Kaluza software generate dot plot masks based on the FSC/SSC parameters and all fluorescence channel combinations to adjust compensation as necessary.
6. Select cells with normal morphology (morpho) using FSC/SSC mask gated on “singlets” (*See 3*.*6*). Then, from the mask FcεR1 FITC/c-kit PE gated in “morpho” generate regions to identify basophils (BASO) as FcεR1^+^c-kit^-^ and mast cells (MC) as FcεR1^+^c-kit and obtain the relative percentage of each gated population by selecting “% of Gated” (*Figure 4*).
7. Obtain histogram plots for CD63 PE and CD200R3 APC by gating on “BASO” to check expression of the two markers in unstimulated and IL-33 stimulated sorted mBaso (*Figure 4*).
8. Generate histogram overlays for CD63 PE to compare expression intensity of this marker on unstimulated and IL-33 stimulated sorted mBaso (*Figure 4*).

**Figure 4.**
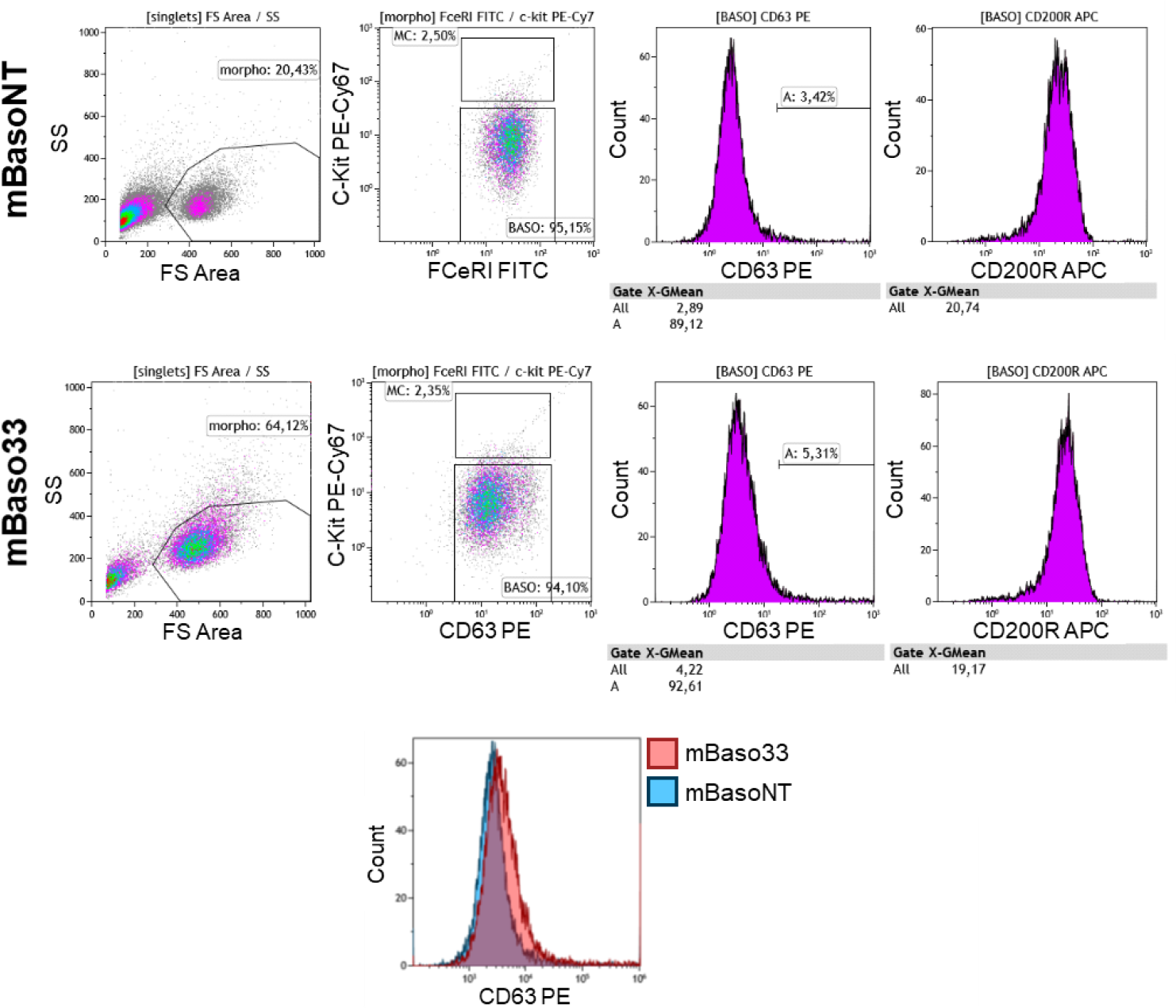
Phenotypic analysis of terminally differentiated and FcεR1+ CD11c- c-kit- sorted mouse basophils, following overnight culture unstimulated or stimulated with IL-33. Following cell sorting, FcεR1+ CD11c- c-kit- mBaso are left unstimulated (mBasoNT) or stimulated overnight with 100 ng/mL rmIL-33 (mBaso33). The following day, cells are stained with the following monoclonal antibodies: FITC anti-mouse FcεR1, PE-Cy7 anti-mouse CD117/c-kit, PE anti-mouse CD63, APC anti-mouse CD200R3. Flow cytometry analysis reveal that, both mBasoNT and mBaso33 maintain basophilic lineage (FcεR1+ CD11c-) and don’t show dedifferentiation in Mast Cells (MC). Moreover, mBaso33 upregulate the activation marker CD63 without down-modulating FcεRI and CD200R3.

## 4) Notes

1. Sacrifice mice and manipulate bones under a Class II Biological Safety Cabinet. BM cells should be cultured for a long lasting differentiation (10 days). For this reason, the entire procedure (starting from tibia and femur collection) requires sterile conditions to avoid bacterial or fungal contamination.
2. ACK red blood cell lysis buffer is prepared with 8.26 g Ammonium Chloride, 1 g Potassium Bicarbonate, 0.037 g EDTA in 1 L of milliq water. Once powders have completely dissolved, the solution is filtered in a 0,22 μm vacuum filter system. ACK is stable for at least 6 months if stored at 4 °C and under ordinary conditions of use.
3. The staining condition is 10-12 minutes in the darkness, +4°C. The antibody dilutions are 1:100 for PE-Cy7 anti-mouse CD117/c-kit, 1:50 for both FITC anti-mouse FcεR1, 1:20 for APC anti-mouse CD200R3 and 1:20 for PE anti-mouse CD49b.
4. Recombinant mouse IL-33 is directly added within cell culture medium. Mix very gently the medium by rotatory movements of the multiwell/flask to ensure a complete dissolution of the cytokine in the culture medium.
5. At the end of overnight culture of mBaso, resuspend the cells by very carefully pipetting before their harvesting. This passage avoids the remaining of mBaso in the well due to their semi-adherent properties.
6. The staining condition is 10-12 minutes in the darkness, +4°C. The antibody dilutions are 1:100 for PE-Cy7 anti-mouse CD117/c-kit and 1:50 for both FITC anti-mouse FcεR1 and APC anti-mouse CD11c.
7. About 2-3×10^5^ mBaso are left unstained and resuspended in 0.5 mL of fresh MACS buffer in a 5 mL polystirene round bottom tube for unstained cell sorting control.
8. This procedure is essential to ensure single-cell suspension and to prevent nozzle clogging.
9. If desired, mBaso can be also stained with a death/viability marker (such as Sytox Blue, 1:1000, directly in MACS buffer) by directly adding it in the ready to sort cell suspension.
10. The staining condition must be performed for 10-12 minutes in the darkness, +4°C. The antibody dilutions are 1:50 for FITC anti-mouse FcεR1, 1:100 for PE-Cy7 anti-mouse CD117/c-kit, 1:20 for PE anti-mouse CD63, 1:20 for APC anti-mouse CD200R3.

## 5) Concluding remarks

A number of findings reported the growing importance of immunomodulatory functions of basophils in health and disease. These rare immune cells exert a potential role in immune responses, particularly Th2 immunity including allergic inflammation and protective roles against parasitic infections (Yamanishi et al., 2017). They are also described to produce a large amount of immunomodulatory cytokines (Seder et al., 1991; Piccinni et al., 1991). Basophils were recently shown to infiltrate human cancers where they can either promote or hamper tumorigenesis (Poto et al., 2022). Due to the crucial and non-redundant roles for basophils, they represent an interesting field of study for immunologists in various disease settings.

The main challenge in the study of this immune cell population, especially in humans, is linked to their scarcity and by the difficult and costly methods for their purification from the blood as well as by ethic issues. A technical issue in human basophil purification is represented by their sensitivity and low purity because of other contaminating circulating cells. In this context, mBaso differentiated *in vitro* may represent an affordable model system to study the immunological functions of these rare immune cells, overcoming limitations in numerical scarcicity and purity of isolated circulating Baso *ex vivo*.

Our methodological contribution provides a standardized procedure to obtain a relevant number of highly purified mouse basophils (FcεR1^+^ CD11c^-^ c-kit^-^) excluding contaminating cells, such as dendritic (CD11c^+^) and mast cell (FcεR1^+^ c-kit^+^). This protocol was succesfully employed for the evaluation of the *in vitro* secretion of cytokines from mBaso as opposed to hBaso in response to alarmins (Gambardella et al., 2022). In conclusion, this methodology allows to obtain a functional population of basophils respecting considerable standards of purity (>98%), without affecting the activation state or viability of these cells.

## Acknowledgments

This work was supported by AIRC (Associazione Italiana Ricerca Cancro) IG 21366 to Giovanna Schiavoni.

## Competing interests

The Authors declare no competing interest.

## Notes

### Competing Interest Statement

The authors have declared no competing interest.

